# Distantly related hepaciviruses share common entry factor, Claudin-1

**DOI:** 10.1101/2022.08.10.503556

**Authors:** Kamilla Toon, Mphatso D Kalemera, Machaela Palor, Nicola J. Rose, Yasuhiro Takeuchi, Joe Grove, Giada Mattiuzzo

## Abstract

Due to increased and broadened screening efforts, the last decade has seen a rapid expansion in the number of viral species classified into the *Hepacivirus* genus. Conserved genetic features of hepaciviruses suggest they have undergone specific adaptation and evolved to hijack similar host proteins for efficient propagation in the liver. Here, we developed pseudotyped viruses to elucidate the entry factors of GB virus-B (GBV-B), the first hepacivirus described in an animal, closely related to hepatitis C virus (HCV). GBV-B pseudotyped viruses (GBVBpp) were shown to be uniquely sensitive to the serum of tamarins infected with GBV-B, validating their usefulness as a surrogate for GBV-B entry studies. We screened GBVBpp infection of hepatoma cell lines CRISPR/Cas9-engineered to ablate expression of individual HCV receptors/entry factors and found that claudin-1 is essential for GBV-B infection, indicating GBV-B and HCV share a common entry factor. Our data suggest that claudin-1 facilitates HCV and GBV-B entry through distinct mechanisms since the former requires the first extracellular loop and the latter is reliant on the second extracellular loop. The implication of this sharing of claudin-1 as an entry factor between these two hepaciviruses on their tissue tropism and evolutionary relationships will be discussed.

**Importance:** Hepatitis C virus (HCV) is a major public health burden; approximately 58 million individuals have chronic HCV infection which could lead to cirrhosis and liver cancer. To achieve the World Health Organisation target of eliminating hepatitis by 2030, new therapeutics and vaccines are needed. Understanding how HCV enters the cells will inform the design of new vaccines and treatments, targeting the first stage of HCV infection. However, the entry mechanism is complex and sparsely described. Studying the entry of related hepaciviruses will increase the knowledge of the molecular mechanisms of the first stages of HCV infection, such as the membrane fusion, and inform structure-guided HCV vaccine design; in this work we have identified a protein, Claudin-1, which facilitates the entry of HCV-related hepacivirus, but with a mechanism not described for HCV. Similar work on other hepaciviruses may unveil commonality of entry factors and possibly new mechanisms.

## Introduction

Hepaciviruses are classified into the Flaviviridae family, which is a broad family of enveloped positive-strand RNA viruses. The *hepacivirus* genus includes hepatitis C virus (HCV), a significant human pathogen affecting up to 3% of the worldwide population, of these, 58 million people are chronically infected [1,2]. GB virus B (GBV-B) was the second hepacivirus described and long remained the only HCV homolog known until another animal hepacivirus was isolated from the nasal swab of a dog with a respiratory illness in 2011 [3]. Since then, exploration of potential animal hosts has uncovered hepaciviral sequences in a diverse range of hosts including dogs, horses, bats, monkeys, rodents, cows, ticks and sharks [4–10]. Studies on these related viruses, including GBV-B, have highlighted the broad hepacivirus spread in the animal kingdom and offered some clues into the origins of HCV [11]. Indeed, characterization of the life cycle of these viruses and the identification of other hepaciviruses could be the key to understand their cross-species transmission and zoonotic potential. Furthermore, animal models of hepaciviral disease could be important surrogate models for HCV disease pathology, and for screening the efficacy of vaccine candidates [12].

GBV-B was isolated from tamarin monkeys that had been experimentally inoculated with the serum of a surgeon (with the initials G.B.) who had presented with the symptoms of acute hepatitis [13–15]. However, the natural host of GBV-B remains enigmatic because of lack of compelling evidence that humans and chimpanzees are susceptible to infection, indicating that the virus, most likely, did not come from the surgeon. In fact, GBV-B has yet to be isolated from any animals other than those experimentally infected [14,16]. Nonetheless, due to HCV’s restricted host tropism, infection of small New World monkeys (tamarins, marmosets, owl monkeys) with GBV-B has previously been employed as a surrogate to study HCV disease and correlates of protections [17–21]. GBV-B, like HCV, is associated with hepatitis and is primarily found in the liver of the infected host [22,23]. However, in contrast to HCV, GBV-B does not appear to cause chronic disease as the infection is usually cleared within six months [16,23–25].

Lack of viral persistence, high costs and the implementation of the “Three Rs” principle in animal research has lessened the GBV-B infection in New World monkey as model for HCV; nevertheless, this model had, and still has, a role in the discovery and evaluation of antiviral and therapeutics for HCV. The tamarin model was key in determining the functional importance of genomic features such as the microRNA-122 binding site [26], and for screening of HCV antivirals such as Ribavirin and NS3 protease inhibitors [27,28]. Furthermore, HCV/GBV-B chimeras developed to overcome the poor sequence identity between the two may yet prove useful in B-cell vaccine candidate screens. These chimeras contain HCV-derived E1E2 glycoproteins (the main target of anti-HCV antibodies), and they can infect marmosets chronically, and show liver pathology consistent with that of HCV in humans [18]. Therefore, marmoset disease progression could be monitored to assess candidate vaccine efficacy.

The hepatotropic nature of most mammalian hepaciviruses described to date suggests these viruses (or common ancestors) have undergone a high degree of evolutionary adaptation to the liver. For instance, the 5’ untranslated regions of most mammalian hepacivirus sequences contain putative binding sites for microRNA-122, which is liver-specific [11]. It can be inferred that hepaciviruses may exploit various orthologous host factors for efficient propagation in the liver. Therefore, HCV’s specific molecular interactions with host factors may be conserved in other hepaciviruses..

A virus, being an obligatory parasite, needs to enter the host cell. HCV exhibits a complex entry mechanism, involving at least four host factors, which is still not fully understood. The virion attaches to hepatocytes via heparan sulfate proteoglycans, at this point, E1E2 glycoproteins engage the scavenger receptor class B type 1 (SR-B1) and the cluster of differentiation 81 molecule (CD81) [29,30]. CD81 engagement is thought to initiate a signalling cascade that leads to intracellular actin remodelling, driving the translocation of the CD81-tethered virus toward the tight junction. HCV then acquires claudin-1 (CLDN1) and occludin (OCLN) during transit to or at the tight junction [31,32]. Finally, the virion is internalized via clathrin-mediated endocytosis and, the process culminates when the low pH environment of the early endosome promotes E1E2-catalysed fusion between the viral and endosomal lipid bilayers [33]. Other entry factors such as the low density lipoprotein (LDLR), epidermal growth factor receptor (EGFR), ephrin receptor A2 (EphA2) and Niemann-Pick C1-like 1 (NPC1L1) have been identified as co-factors for entry [33–36].

Recently, SR-B1 has been shown to mediate the entry of a cell culture derived rat hepacivirus [37]. Aside from this and the HCV entry factors outlined above, no other entry factors have been identified for the other hepaciviruses. *In vitro* studies have been impaired by the scarcity of replication-competent full-length cell culture viruses. However, the discovery of entry factors could reveal mechanistic details surrounding HCV entry, thus potentially informing vaccine design. Moreover, such studies could uncover a novel mechanism of lipid bilayer fusion as hepaciviral E1E2 are genetically and structurally predicted to belong to a novel class of membrane fusion protein, outside of the three described so far [38]. In this report, we adapted the well-established HCV pseudotyped virus (HCVpp) system to study GBV-B cell entry. HCVpp are retroviral-based particles bearing HCV E1E2 glycoproteins. HCVpp were crucial in the identification of CLDN1 and OCLN as HCV entry factors [39–41], as well as in the determination of epitopes that are targeted by neutralizing antibodies during infection [42–44]. Here, we generated GBV-B pseudotyped virus (GBVBpp) and then employed a receptor knock-out cell-line screen to characterize GBV-B entry. We found that, like HCV, GBV-B entry is dependent on CLDN1, but that the genetic determinants of this vary between the viruses, suggesting subtle mechanistic differences.

## Results

### Production of GBV-B Pseudotyped virus

To study and compare the entry mechanisms between GBV-B and HCV, a retroviral vector based on murine leukaemia virus (MLV), encoding a firefly luciferase reporter gene, was produced similar to previous work [21]. Infectivity of the HCV pseudotype (HCVpp) and GBV-B pseudotype (GBVBpp) was assessed on an immortalised human cell line Huh7.5 and detected as luminescence signal (Figure 1A). Infectivity over background at similar level as HCVpp could only be obtained when GBVBpp were produced using codon optimized E1E2 sequences, which was therefore chosen for the production of GBVBpp. These pseudotyped particles were then tested against archived serum from a tamarin experimentally infected with GBV-B [17] (Figure 1B). Infectivity of GBVBpp was inhibited in a dose dependant manner by serum from GVB-B infected tamarins, but not by serum from a naïve tamarin (Figure 1C); this is consistent with specific entry driven by GBV-B E1E2 and indicates that GBVBpp represents a suitable model for entry and serological studies.

**Figure 1.**
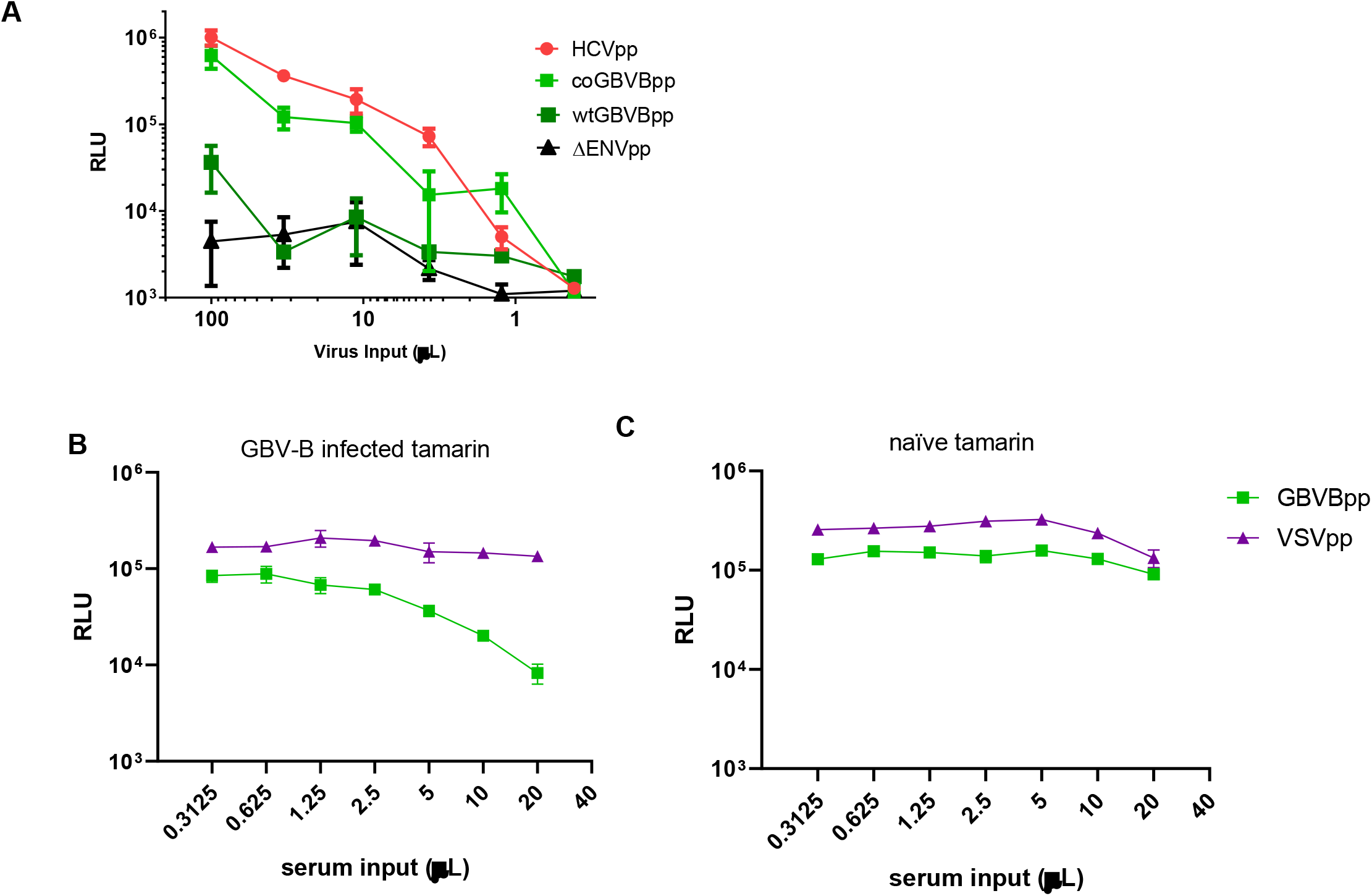
Production of MLV core pseudotyped with GBV-B E1E2. **(A)** HCVpp (isolate UKN1A20.8), GBVBpp, which was produced using a plasmid with GBV-B E1E2 sequences cloned from ATCC stock (wtGBVBpp) or codon optimized for mammalian expression (coGBVBpp), and ΔENVpp were produced with an MLV vector encoding a luciferase reporter gene. Producer cell supernatant containing particles were titrated on Huh 7.5 cells. **(B-C)** GBVBpp, with the codon optimized E1E2 sequence, were incubated with increasing volumes of serum from a tamarin experimentally infected with GBV-B (**B**) or from a naïve animal (**C**) prior to infect Huh7.5 cells. MLV vector pseudotyped with the vesicular stomatitis virus glycoprotein G (VSVpp) was also tested against the same tamarin sera to confirm specificity of neutralization. Infectivity is expressed as luciferase activity in relative luminescence units (RLU). RLU values are mean +/- standard deviation of one experiment in triplicate. These are representative of 2 independent experiments.

### CLDN-1 mediates GBV-B entry in human cells

Given that GBV-B is closely related to HCV and is also hepatotropic, we hypothesized that the two viruses may share conserved entry factors. To this end, a panel of HCV receptor knockout (KO) Huh7 cell lines [45] was screened for susceptibility to GBVBpp entry (Figure 2A). No effect was observed in CD81, OCLN, LDLR, or SR-B1 KO cell lines; however, GBVBpp entry was significantly reduced in CLDN1 KO cells (Fig 2A). An unrelated pseudotyped MLV carrying the vesicular stomatitis virus glycoprotein G (VSVpp) was used as negative control and infected CLDN1 KO cells as efficiently as unmodified Huh7 cells.

**Figure 2.**
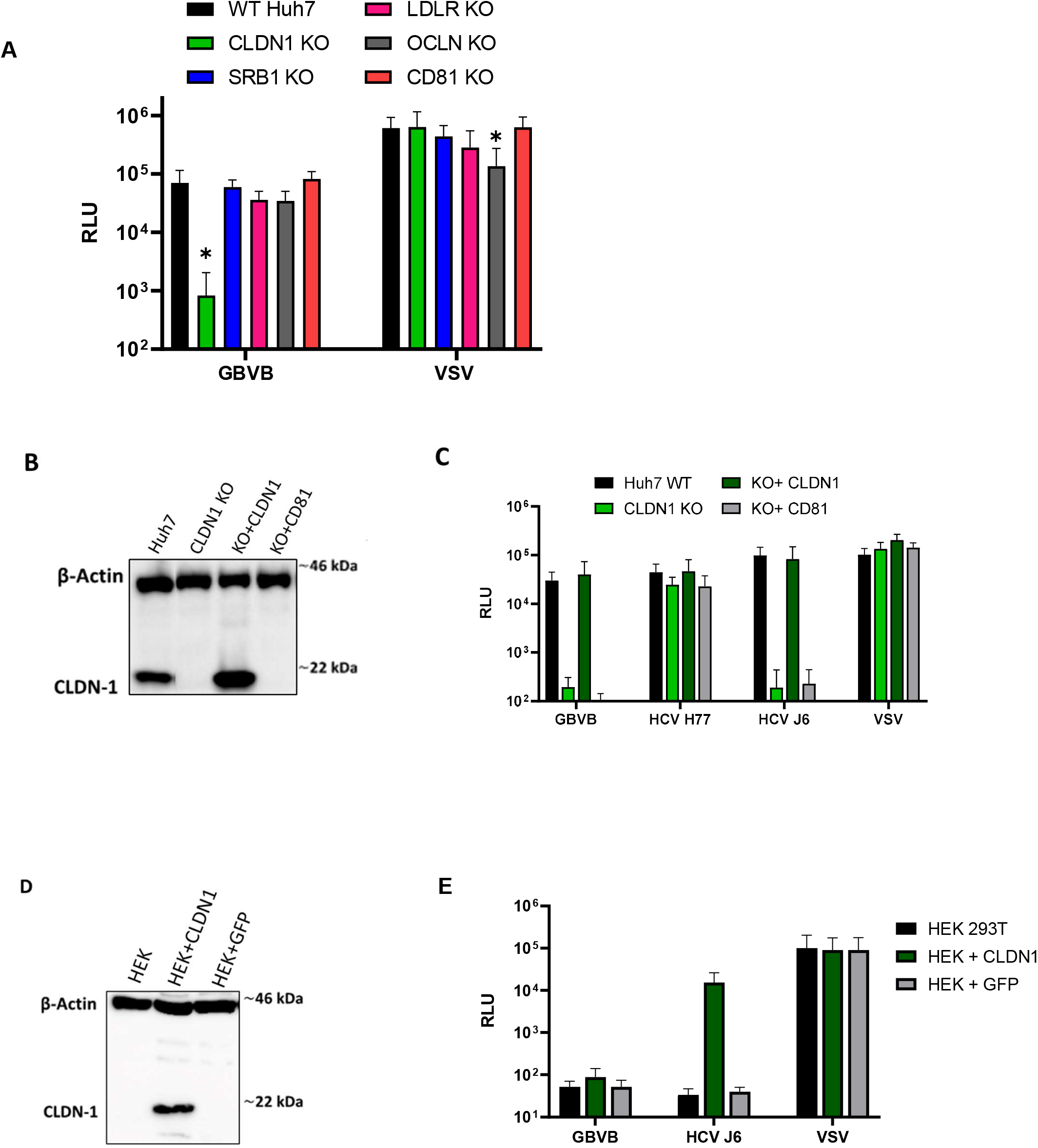
CLDN-1 is required for GBV-B entry in human cells. **(A)** Huh7 cells in which CLDN1, SR-B1, LDLR, OCLN, or CD81 were knocked out were infected with GBVBpp or VSVpp. **(B-D)** CLDN1 expression in modified Huh7 **(B)** or HEK 293T (**D**) cells was assessed by immunoblotting using an anti-CLDN1 antibody. Protein input was verified using an anti-actin antibody. Samples were run with a protein size marker; size in kilo Dalton (kDa) is indicated to the right of the blot. The blots are consistent with the predicted molecular weight of CLDN1 being 23 kDa. **(C)** Huh7 cells, Huh7 CLDN1 KO cells transduced with lentiviral vectors to express CLDN1 or CD81 were challenged with the indicated PVs. **(E)** HEK 293T cells were transduced to express CLDN1 or and irrelevant protein (GFP) and challenged with GBVB, HCV, or VSV pseudotyped virus. Infectivity is expressed as mean RLU values +/- standard deviation of 2 independent experiments run in triplicate *t-test p<0.01.

To further confirm the role of CLDN1 in GBV-B entry, CLDN1 expression in KO cells was reconstituted to determine if this restored susceptibility to GBVBpp. CLDN1 KO cells were transduced with a lentiviral vector encoding CLDN1 or an irrelevant gene (CD81) (Figure 2B). The exogenous expression of CLDN1, but not CD81, restored susceptibility of KO cells to GBVBpp and HCV J6pp, a CLDN1-restricted strain of HCV, to wildtype levels (Figure 2C). This result further supports that CLDN1 is necessary for GBV-B entry. Also, expression of CLDN1 above endogenous levels does not enhance entry, consistent with what has been observed for HCV entry [31]. HCVpp carrying isolate H77-derived E1E2 could similarly infect CLDN1 KO and wildtype cells, consistent with previous findings that this isolate can use CLDN6 or CLDN9 as well as CLDN1 for entry.

HEK293T (HEK) cells are non-permissive to HCV entry as they do not express CLDN1. However, upon exogenous expression of CLDN1, HEK cells become highly permissive to HCVpp, indicating that all other HCV entry factors are present [30]. CLDN1-transduced HEK cells (Figure 2D) were susceptible to HCVpp infection, whereas no signal was detected for GBVBpp (Fig 2E). This observation suggests that there is at least one other entry factor present in Huh7, but not HEK cells, that is essential for GBV-B entry, consistent with its liver tropism. Alternatively, there may be unknown inhibitory factors in HEK cells specific to GBV-B, but not to HCV.

### CLDN1 critical regions for GBV-B entry differs from those required for HCV

Claudin-1 is a small protein (211 aa) located at the cell surface with 4 transmembrane domains and 2 extracellular loops, with the N- and C-termini both intracellular [46]. Previous work has demonstrated that amino acid residues at positions 32 and 48 in the first extracellular loop (EL1) of CLDN1 are crucial to support HCV entry [31]. We sought to determine whether the same residues are similarly important for GBV-B infection. Through site-directed mutagenesis, we generated a lentiviral vector for an CLDN1 mutant expressing isoleucine to methionine at position 32 (I32M) and glutamic acid to lysine at position 48 (E48K) substitutions in CLDN1 EL1 (Figure 3A). I32M/E48K mutant was then introduced into CLDN1 KO cells through lentiviral transduction and confirmed by western blot (Figure 3B). The mutant, as expected, failed to rescue HCV J6pp entry (Figure 3C). Notably, mutant and wildtype CLDN1 restored GBVBpp infectivity to a similar degree (Figure 3C), indicating I32 and E48 of CLDN1 are not essential for GBV-B’s cell entry.

**Figure 3.**
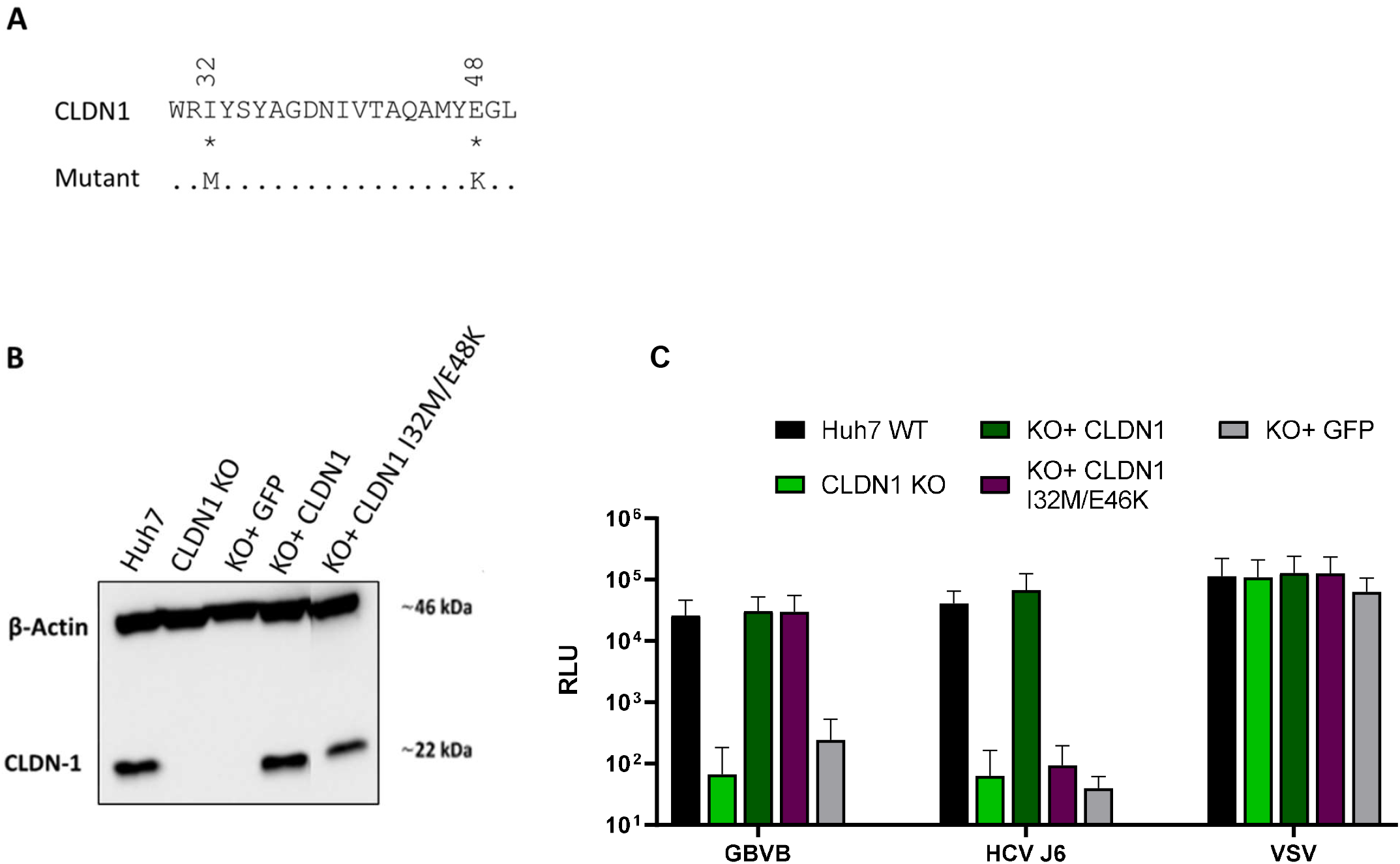
Different CLDN1 regions are important for GBVB and HCV’s entry. **A)** Alignment of CLDN1 amino acids 30-50 in extracellular loop 1 with the mutant created by site directed mutagenesis to introduce I32M and E48K. Identical amino acids are represented by a full stop and numbering represents the amino acid position in the full length CLDN1 (NP_066924.1). (B) CLDN1 expression in modified Huh7 cells was assessed by immunoblotting using an anti-CLDN1 antibody. Protein input was verified using an anti-actin antibody. Samples were run with a protein size marker; size in kDa is indicated to the right of the blot. (C) Huh7 CLDN1 KO cells were transduced to express CLDN1, CLDN1 I32M/E48K, or GFP only and challenged with MLV-based pp harbouring the indicated envelope proteins. Infectivity is expressed as mean RLU values +/- standard deviation of 2 independent experiments run in triplicate. Infectivity is expressed as mean RLU values +/- standard deviation of 2 independent experiments run in triplicate.

### Characterization of CLDN1 regions important for GBV-B entry

We next sought to determine which region of CLDN1 is important for GBV-B entry through domain swapping studies. CLDN6 and CLDN9 were selected as non-permissive candidates; this choice was driven by the observation that CLDN1-independent HCV strain H77 can enter CLDN1 KO cells (Figure 2B) which indicates that CLDN6 and/or CLDN9 are expressed in Huh7 cells, but GBVBpp is unable to utilize them for cell entry. CLDN6 and CLDN9 genes were sub-cloned from Huh7 cells into a lentiviral vector and used to transduce CLDN1 KO cells. Only the addition of CLDN1, and not overexpression of CLDN6 or CLDN9, conferred GBVBpp and HCV J6pp susceptibility of KO cells (Figure 4A). HCV H77pp infection of CLDN6 and CLDN9 transduced HEK cells confirmed ectopic protein was correctly folded and functional (Figure 4B). CLDN9 was chosen for domain swapping experiments as it has slightly higher homology to CLDN1 at the amino acid level than CLDN6, 45% versus 43%. An overlapping PCR strategy was employed to generate CLDN1/CLDN9 chimeras reciprocally swapped for their extracellular loops (EL1 and EL2) (Figure 4C).

**Figure 1.**
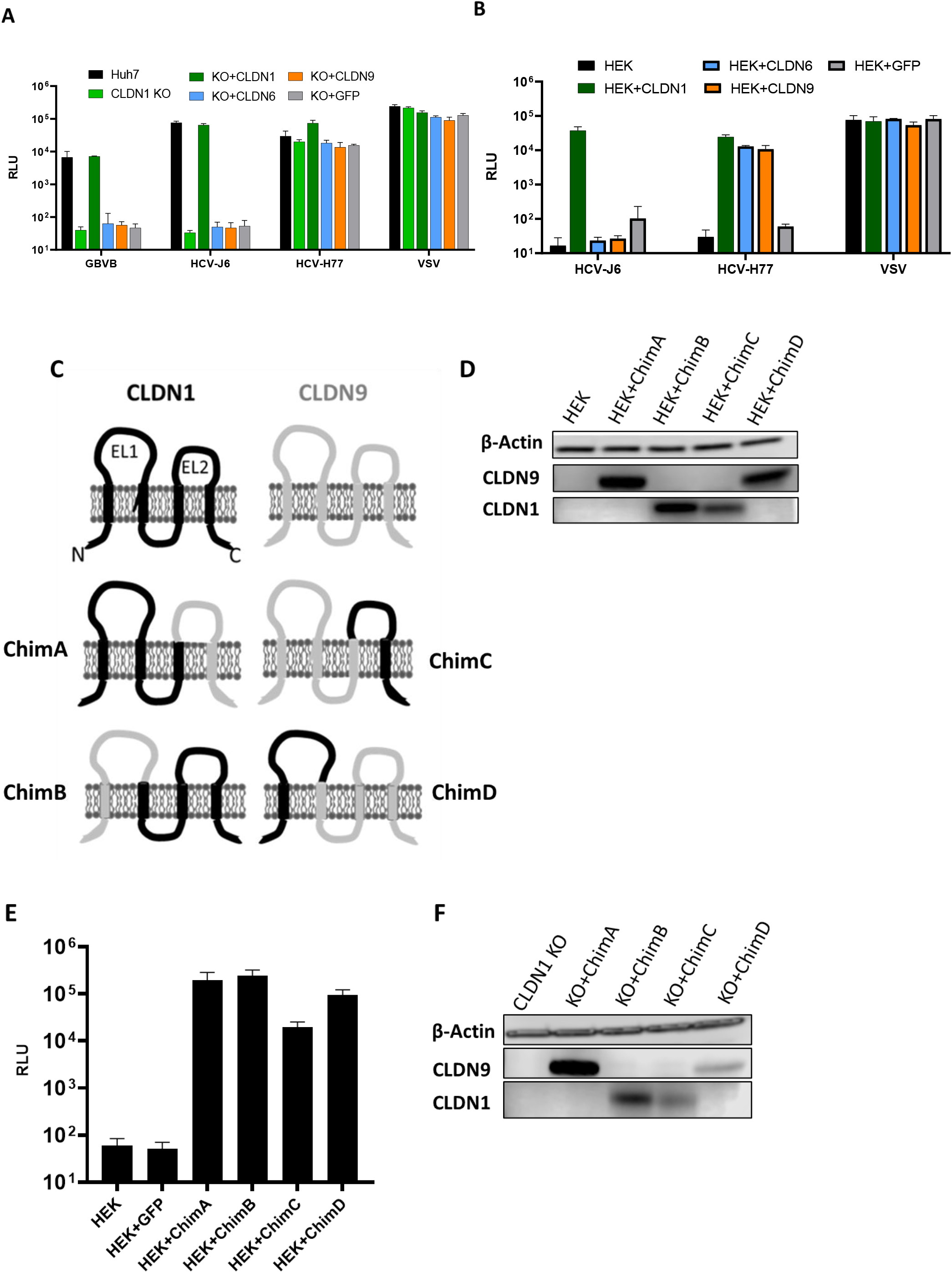
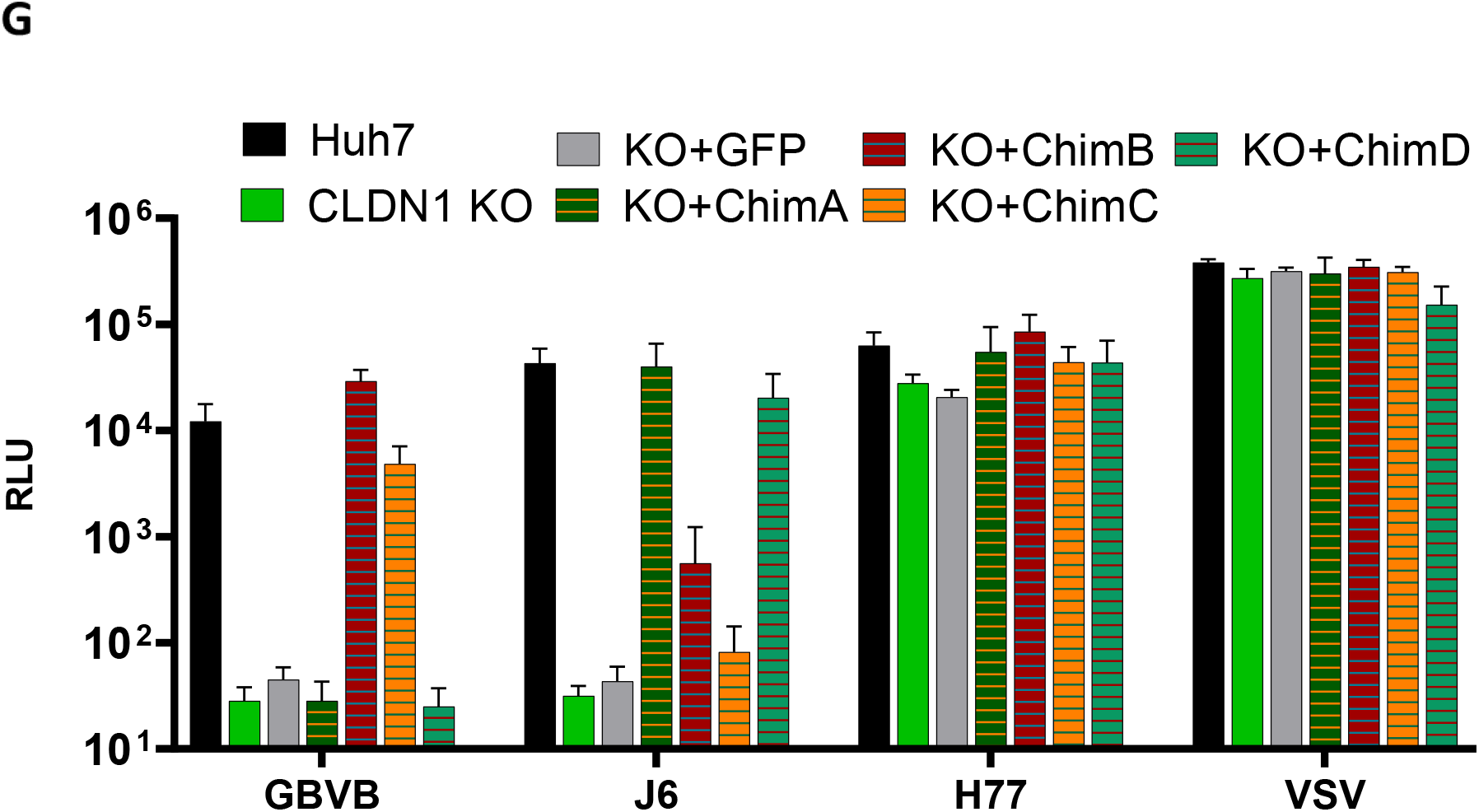
Extracellular loop 2 of Claudin-1 is important for GBVB entry. Huh7 CLDN1 KO **(A)** or HEK 293T cells **(B)** transduced to express CLDN1, CLDN9, CLDN6 or GFP proteins were challenged with pp harbouring glycoproteins from GBVB, HCV J6, HCV H77 and VSV. **(C)** Claudin-1 (black) and Claudin-9 (grey) topology with the indicated regions of each protein swapped with respective region of the other CLDN to produce chimeric proteins, resulting in chimeric proteins with the following amino acids: ChimA (CLDN1 1-139/CLDN9 138-217), ChimB (CLDN9 1-81/CLDN1 82-211), ChimC (CLDN9 1-137/CLDN1 139-211), and ChimD (CLDN1 1-81CLDN9 82-217). Chimeric protein expression in HEK cells **(D)** or Huh7 cells **(F)** was assessed by immunoblotting using anti-CLDN1 and anti-CLDN9 antibodies. Protein input was verified using an anti-actin antibody. HEK 293T cells transduced to express the indicated chimeric CLDN1/9 were infected with H77pp **(E)**. Huh7 CLDN1 KO cells transduced to express the indicated chimeric CLDN1/9 were infected with GBVBpp J6pp, J77pp and VSVpp (G). Infectivity is expressed as mean RLU values ± standard deviation of 2 independent experiments run in triplicate.

The chimeric CLDN1/9 proteins were initially transduced into HEK cells and expression confirmed by western blot (Figure 4D). The anti-CLDN1 and CLDN9 antibodies appear to recognize a region in the C-terminus of their respective protein; therefore, ChimA and ChimD are detected by anti-CLDN9 antibodies and ChimC and ChimB are detected by anti-CLDN1 antibodies. To assess functionality of these chimeric CLDN1/9 proteins, transduced HEK cells were challenged with HCV H77pp, which can use both CLDN1 and CLDN9. Figure 4E shows the complete panel of chimeric CLDN1/9 proteins expressed in HEK cells, which have conferred susceptibility to H77pp while the parental cells and the transduction control (expressing GFP only) remain uninfectable by the virus. This confirms that all chimeras are conformationally correct and support HCV infection.

The panel of chimeras was then expressed in Huh7 CLDN1 KO cells (Figure 4F) and tested for susceptibility to GBVBpp and J6pp (Figure 4G). We found that GBVBpp could infect CLDN1 with EL1 replaced with that of CLDN9 (ChimB), but infection fell to below the limit of detection when EL2 is swapped (ChimA); suggesting that the EL2 of CLDN1 is critical for GBV-B entry. Consistently, GBVBpp infection was observed when EL2 of CLDN1 was introduced to CLDN9 (ChimC). Furthermore, no infectivity is observed when EL1 is introduced (ChimD), confirming that EL2, or a region downstream, is the critical region. The opposite was observed for HCV J6pp, as expected as HCV is known to utilize EL1 [31]. The difference in CLDN-1 regions necessary for GBV-B and HCV infection suggests a novel way of interaction with CLDN-1 than that described for HCV.

## Discussion

Much of what is known of the hepaciviral life cycle is inferred from HCV, which is by far the most extensively studied virus in the genus due to the significant disease burden it poses to humans. GBV-B was sequenced in 1995 and since has been a useful model for HCV research [18,26–28]; despite this, no receptors or entry factors had been described for it, until now. Here, we demonstrate that like HCV, CLDN1 is an entry factor that is necessary for GBV-B cell entry. When CLDN1 is knocked out of Huh7, a susceptible cell line to GBV-Bpp, entry is diminished (Figure 2A) and upon exogenous expression of CLDN1 susceptibility is completely restored (Figure 2B).

In addition to HCV, coxsackievirus B, some reoviruses as well as some adenoviruses also utilize tight junction proteins as coreceptors [47,48]. In polarized cells, the majority of CLDN1 is localized in tight junctions [31]. HCV is thought to follow a similar cell entry pathway as coxsackievirus B, where it binds a primary receptor on the luminal cell surface then migrates laterally along the plasma membrane to encounter tight junction proteins CLDN1 and OCLN [33]. However, there is a small amount of CLDN1 that localize at the basal membranes of hepatocytes, and there is evidence to support that HCV can utilise this fraction of CLDN1 [49–51]. Of note, GBVBpp were insensitive to OCLN deletion which may further highlight a divergence in the entry pathways of HCV and GBV-B since OCLN, which exclusively localizes at the tight junctions, is indispensable for HCV infection.

Two amino acids in the extracellular loop 1 (EL1) of CLDN1, previously identified as essential for HCV entry [31] have no impact on GBV-Bpp infection (Fig 3B). These residues are responsible for the interaction of CLDN1 with CD81 to form a complex that is indispensable for HCV entry [52]. It is not surprising that these residues are not important for GBV-B entry because CD81 does not appear to play a role in GBV-B entry into human cells (Figure 2A); thus, inability to form this complex was expected to have no effect, further confirming that GBV-B entry is independent of CD81. We have also shown that GBV-B is dependent on the EL2, or a downstream region of CLDN1 for infection. Using chimeric proteins between permissive CLDN1 and nonpermissive CLDN9, GBVBpp were able to enter only those cells expressing a chimeric CLDN containing CLDN1 EL2, while CLDN1-dependent HCV strains J6pp were able to infect cells expressing chimeras with EL1 of CLDN1 (Figure 4G). Further investigation is needed to determine the specific residues of EL2 that are important for GBV-B entry and whether GBV-B directly interacts with CLDN1.

It was initially thought that CLDN1 does not directly interact with HCV particles but is important for the receptor complex formed with CD81 [52,53]. However, some studies have indicated there may be direct interaction between CLDN1 and HCV E1E2 complex. CLDN1 has been shown to not interact with E2 alone, but with the E1E2 heterodimer [54], and mutations in E1 have been shown to shift use of CLDN1 to CLDN6 [55,56], possibly indicating a direct interaction with E1. However, this interaction is poorly understood; it is not known which domains on CLDN1 are important for interaction with the glycoproteins or if binding to CLDN1 is needed for receptor clustering. It is also currently unknown whether CLDN1 binds with E1 or whether E1E2, together, form a conformational domain for interaction.

Another discrepancy observed between HCVpp and GBVBpp is the ability to infect HEK cells transduced to expressed CLDN1: these are susceptible to HCVpp but not GBVBpp (Figure 3E). This suggests that GBV-B may utilize at least one entry factor not conserved between GBV-B and HCV that is expressed in Huh7 cells, but not in HEK293 cells. Thus, further investigation into other entry factors is needed to fully characterize GBV-B cell entry, for example, through a CRISPR screen of liver-enriched cell membrane proteins. Identification of further receptors or entry factors that are not conserved may shed light on the physiological differences seen between the new world monkey animal model for GBVB and HCV infection. Alternatively, HEK293, but not Huh7, cells may have a factor restricting GBV-B entry.

A caveat to our investigations is that GBVBpp may not be representative of authentic replicating viral particles. Being closely related with a similar hepatotropism, it is likely that GBV-B particles, like HCV, resemble low-density lipoprotein complexes associated with apolipoproteins [57,58]; therefore, apolipoprotein receptors such as LDL-R and SR-BI are likely to play significant role in virus entry as they can tether apolipoprotein-associated virions to the basolateral surface of hepatocytes [30,34,45]. Indeed, it was recently shown by Scheel and colleagues that an infectious cell culture-derived rat hepacivirus also shared biophysical properties with LDL and was dependent on SR-B1 for entry [37]. The development of culture-derived replication-competent GBV-B *in vitro* and subsequent ultrastructural analyses of this virus could help establish whether co-option of the LDL biogenesis pathway is a conserved feature among hepaciviruses.

GBV-B provides a potential avenue to investigate the interaction between E1E2 glycoproteins and CLDN1 without reliance on the CD81 complex, potentially helping further define CLDN1’s role in HCV cell entry. Characterisation of the entry mechanism may unveil new potential targets for therapeutics and uncover a novel mechanism of membrane fusion. Unlike the flavivirus entry protein E, hepaciviral E1E2 proteins are not thought to be class II membrane fusion proteins; indeed, structural analyses suggest that hepacivirus E1E2 may represent a novel class of fusion protein [59–62]. To this end, studying GBV-B E1E2 entry may prove the more facile path to deciphering the hepaciviral membrane fusion mechanism as it is structurally predicted to have fewer disordered segments compared to HCV. Solving the hepaciviral fusion mechanism could inform structure-guided HCV vaccine design. Understanding the similarities and differences between HCV and GBV-B E1E2 will likely provide mechanistic insight into HCV’s complex entry and possibly offer clues as to why it seems HCV has uniquely evolved to cause chronic disease in humans, while close relatives in other hosts succumb to immune pressure in the acute stages of infection.

In summary, we have discovered that two distantly related hepaciviruses, HCV and GBV-B, share CLDN-1 as a cell entry factor, but their mode of CLDN1 usage is different. This may suggest that this shared factor usage may have arisen by convergence through two different evolutionary routes. Alternatively, use of CLDN1 as an entry factor by a hepacivirus ancestor along with the establishment of liver tropism may have been retained with certain differentiation of interaction mode by divergent extant hepaciviruses. We expect future studies on other animal hepacivirus cell entry will shed some light on these evolutionary relationships between hepaciviruses and cellular entry factors.

## Materials and Methods

### Cells

HEK 293T cells were obtained from the American Tissue Culture Collection (CRL-11268). Huh7 and HCV receptor knockout cell lines were kindly supplied by Prof Yoshiharu Matsuura (Osaka University) [45]. To genetically modify cell lines, 1 million Huh7 CLDN1 KO or HEK 293T cells were seeded in 6 well plates and approximately 2 hours later were transduced with VSV-G pseudotyped lentiviral vectors expressing GFP and the protein of interest at MOI of approximately 1, in the presence of 8µg/ml of polybrene. All cells were maintained at 37°C, 5% CO_2_ in Dulbecco’s Modified Essential Medium (DMEM) supplemented with Glutamax (Gibco), 10% foetal bovine serum (Pan Biotech), 100 unit/mL penicillin, 100 μg/mL streptomycin (Sigma), and 1% non-essential amino acids (Gibco).

### Tamarin sera

Archived sera from GBV-B infected red-bellied tamarins (*Saguinus labiatus*) were available from a previous study [17].

### Pseudotyped Virus Production

To produce lentiviral or gammaretroviral pseudotyped virus, 4 million HEK 293T cells were seeded in a 10cm dish to reach 50-70% confluence. The day after, a DNA plasmid mix was prepared containing 1μg of plasmid encoding either MLV or HIV structural and enzymatic proteins, pCMVi [63] or p8.91 [64] respectively, 1μg of viral envelope protein expression plasmid and 1.5μg of transfer vector plasmid expressing a luciferase (pCFCR-LUC) or GFP reporter gene (pDual) in 15µl of Tris-EDTA buffer. The DNA plasmid mix was added to 18µl of Fugene-6 (Promega) transfection reagent in 200µl of pre-warmed Opti-MEM (Gibco) and incubated for 20 minutes. The transfection mix was then added to the cells which were incubated at 37°C, 5% CO_2_ for 24 hours before media was replaced. After 48 hours, supernatant was collected and filtered through 0.45µm cellulose acetate membrane (BioWhittacker). Pseudotyped viral particles contained in the supernatant were concentrated by either 1240 RCF, overnight at 4°C, or 103,586 RCF, for 2 hours at 4°C.

### Plasmids

Full length GBV-B E1E2, nucleotides 851-2449 (NC_001655.1), and codon optimized E1E2 genes (synthesized by Genewiz) were cloned into pCAGGS [65]. pD607_J6_E1E2 [66] and pD603_H77_E1E2 (Addgene # 86983) are mammalian expression vectors encoding the codon optimized HCV glycoproteins from strains J6 and H77, respectively. pDual_CLDN1 or pDUAL_CD81 are lentiviral dual promoter transfer vectors expressing GFP and CLDN1 or CD81, respectively (Addgene #86981, #86980). Mutant (I32M and E48K) pDual_CLDN1 was created using Q5® site directed mutagenesis kit (New England Biolabs) with the following primers I32M_F (5’-CCA GTG GAG GAT GTA CTC CTA TGC C-3’), I32M_R (5’-GGC AGG GCA GTG CTG ACG-3’), E48K_F (5’-GGC CAT GTA CAA GGG GCT GTG GA-3’), and E48K_R (5’-TGG GCG GTC ACG ATG TTG-3’). To clone CLDN6 and CLDN9 genes, RNA from Huh7 cells was extracted with RNeasy kit (Qiagen) according to the manufacturer’s instructions. cDNA was then synthesized from the RNA with Superscript IV transcriptase kit (Invitrogen) according to manufacturer’s instructions using the following primers CLDN6_F (5’-AAT TAG GAT CCG CCG CCA CCA TGG CCT CTG CCG GAA TGC A-3’), CLDN6_R (5’-GCG GCG GCC GTC GAC TCA GAC GTA ATT CTT GGT AGG GTA-3’), CLDN9_F (5’-AAT TAG GAT CCG CCG CCA CCA TGG CTT CGA CCG GCT TAG A-3’), and CLDN9_R (5’-GCG GCG GTC GAC TCA CAC GTA GTC CCT CTT-3’). Amplified genes were subcloned in the pDual lentiviral vector using *BamHI* restriction site at the 5’ and a *SalI* restriction site at the 3’ end of the gene, introduced with the primers. Successful cloning was confirmed through Sanger sequencing.

### Chimeric protein production

Chimeric proteins containing the sequences of CLDN1 and CLDN9 proteins were spliced using PCR driven overlap extension as described in [67]. Briefly, a first PCR was performed using the templates and primers shown in Table 1 with KOD hot start DNA polymerase (Sigma-Aldrich), with annealing time of 30 seconds at the indicated temperature. Primers A and B or C and D (Table 1) were designed to introduce an overlapping sequence of 10-12 nucleotides to each protein fragment that spans the junction where the proteins will be spliced together. In a second PCR, the overlapping sequences created by the first PCR were annealed together to join the sequences of CLDN1 and CLDN9, and primers A and D were used to amplify the hybridized product. Correct product synthesis was verified by Sanger sequencing.

**Table 1.**
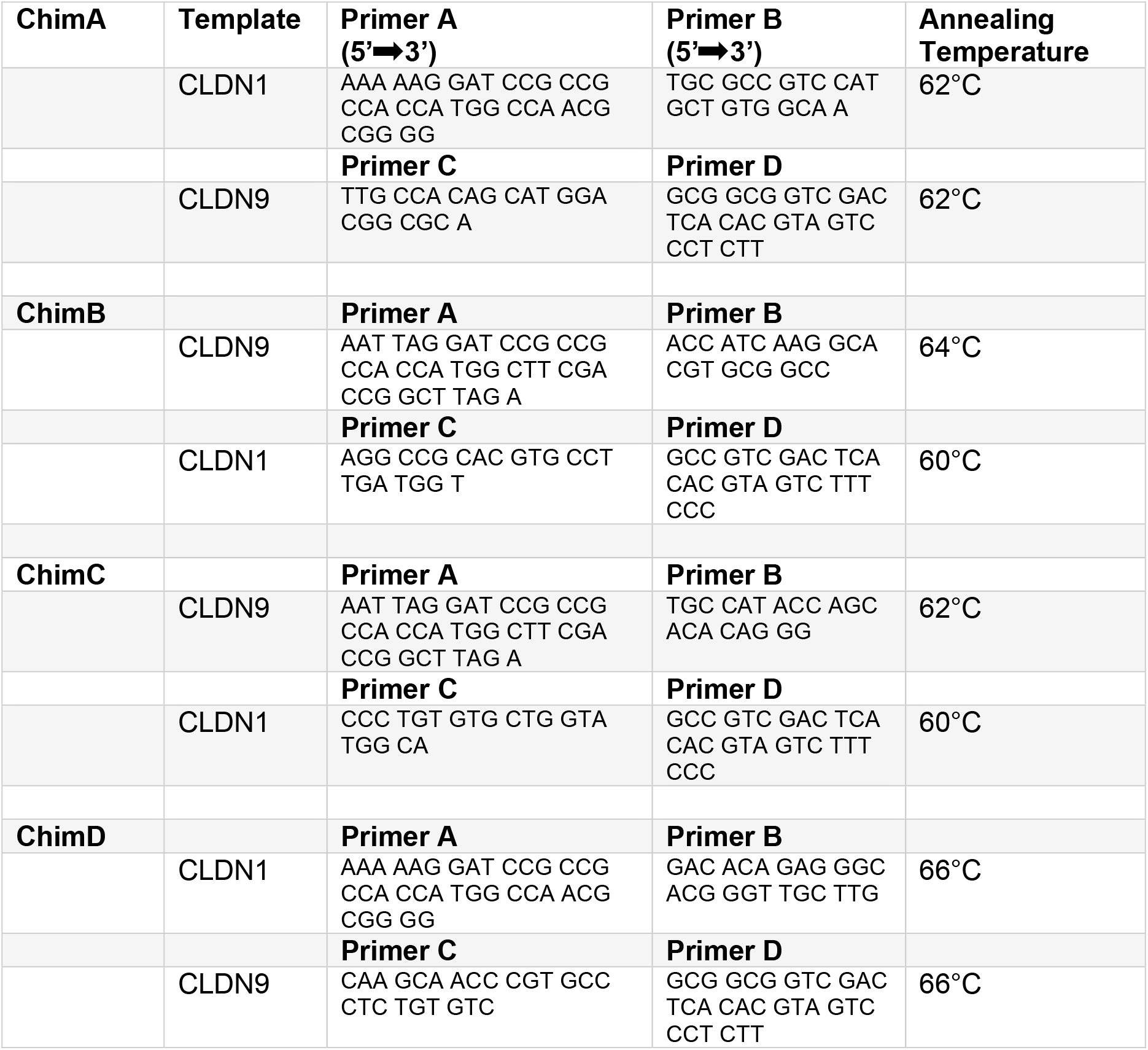
Overlap extension PCR primers.

### Immunoblotting

Cells resuspended in lysis buffer on ice for 5 minutes as previously described [68], centrifuged for 5 mins and the supernatant collected for analysis. Proteins were separated by SDS-PAGE in a 4-20% tris-glycine gel. Proteins were transferred to a nitrocellulose membrane and non-specific binding blocked by incubation with 2 % milk, 0.1 % Tween-20 in PBS. Membranes were probed by serial incubation with rabbit anti-CLDN1 antibody (1:1000 Abcam) or rabbit anti-CLDN9 antibody (1:300 Proteintech), rabbit anti-βactin antibody (1:10000 Abcam), and goat anti-rabbit secondary conjugated to horseradish peroxidase (HRP). Chemiluminescence signal was then measured using Chemidoc MP (Bio Rad) or D-Digit blot scanner (LI-COR).

### Infectivity Assays

For pseudotyped particles with a luciferase reporter, target cells were seeded at 15,000 cells per well in a 96-well plate and infected with supernatant containing pseudotyped particles, with a final concentration of 4 µg/mL of polybrene. Plates were spin-inoculated at 1240 RCF for 30 min, at 20°C. Approximately 72 hours later, cells were lysed by adding 100 µl of 1:1 (v:v) mixture of phenol red-free DMEM and Bright-Glo substrate (Promega) and luminescence detected using a Glomax Navigator (Promega). For pseudotypes carrying a GFP reporter gene, target cells were seeded at 100,000 per well in a 24-well plate and transduced with supernatant containing pseudotypes, with a final concentration of 4µg/mL of polybrene and incubated at 37°C for 72 hours. Cells were then detached from plate with trypsin, fixed with 4% paraformaldehyde in PBS, and analysed on a FACS CANTO II for GPF expression.

## Acknowledgments

We thank Prof Yoshiharu Matsuura (Research Institute for Microbial Diseases, Osaka University, Osaka, Japan) for kindly proving the HCV receptor knockout cells. K.T. was supported by a PhD studentship through the National Institute for Biological Standards and Control, the Medicines and Healthcare products Regulatory Agency. J.G. is supported by a Sir Henry Dale fellowship from the Wellcome Trust and Royal Society (107653/Z/15/A) and by the Medical Research Council (MC_UU_12014).

## References

1. World Health Organization, Hepatitis C Available online: www.who.int/news-room/fact-sheets/detail/hepatitis-c (accessed on 20 July 2022).

2. Chigbu, D.; Loonawat, R.; Sehgal, M.; Patel, D.; Jain, P. Hepatitis C Virus Infection: Host–Virus Interaction and Mechanisms of Viral Persistence. Cells 2019, 8, 376, doi:10.3390/cells8040376.

3. Kapoor, A.; Simmonds, P.; Gerold, G.; Qaisar, N.; Jain, K.; Henriquez, J.A.; Firth, C.; Hirschberg, D.L.; Rice, C.M.; Shields, S.; et al. Characterization of a Canine Homolog of Hepatitis C Virus. Proc. Natl. Acad. Sci. 2011, doi:10.1073/pnas.1101794108.

4. Pfaender, S.; Cavalleri, J.M.V.; Walter, S.; Doerrbecker, J.; Campana, B.; Brown, R.J.P.; Burbelo, P.D.; Postel, A.; Hahn, K.; Anggakusuma; et al. Clinical Course of Infection and Viral Tissue Tropism of Hepatitis C Virus-like Nonprimate Hepaciviruses in Horses. Hepatology 2015, 61, 447–459, doi:10.1002/hep.27440.

5. Kapoor, A.; Simmonds, P.; Scheel, T.K.H.; Hjelle, B.; Cullen, J.M.; Burbelo, P.D.; Chauhan, L. V.; Duraisamy, R.; Sanchez Leon, M.; Jain, K.; et al. Identification of Rodent Homologs of Hepatitis C Virus and Pegiviruses. MBio 2013, doi:10.1128/mBio.00216-13.

6. Quan, P.-L.; Firth, C.; Conte, J.M.; Williams, S.H.; Zambrana-Torrelio, C.M.; Anthony, S.J.; Ellison, J.A.; Gilbert, A.T.; Kuzmin, I. V.; Niezgoda, M.; et al. Bats Are a Major Natural Reservoir for Hepaciviruses and Pegiviruses. Proc. Natl. Acad. Sci. 2013, 110, 8194–8199, doi:10.1073/pnas.1303037110.

7. Lauck, M.; Sibley, S.D.; Lara, J.; Purdy, M.A.; Khudyakov, Y.; Hyeroba, D.; Tumukunde, A.; Weny, G.; Switzer, W.M.; Chapman, C.A.; et al. A Novel Hepacivirus with an Unusually Long and Intrinsically Disordered NS5A Protein in a Wild Old World Primate. J. Virol. 2013, 87, 8971, doi:10.1128/JVI.00888-13.

8. Corman, V.M.; Grundhoff, A.; Baechlein, C.; Fischer, N.; Gmyl, A.; Wollny, R.; Dei, D.; Ritz, D.; Binger, T.; Adankwah, E.; et al. Highly Divergent Hepaciviruses from African Cattle. J. Virol. 2015, 89, 5876, doi:10.1128/JVI.00393-15.

9. Shi, M.; Lin, X.-D.; Vasilakis, N.; Tian, J.-H.; Li, C.-X.; Chen, L.-J.; Eastwood, G.; Diao, X.-N.; Chen, M.-H.; Chen, X.; et al. Divergent Viruses Discovered in Arthropods and Vertebrates Revise the Evolutionary History of the Flaviviridae and Related Viruses. J. Virol. 2016, 90, 659, doi:10.1128/JVI.02036-15.

10. Harvey, E.; Rose, K.; Eden, J.-S.; Lo, N.; Abeyasuriya, T.; Shi, M.; Doggett, S.L.; Holmes, E.C. Extensive Diversity of RNA Viruses in Australian Ticks. J. Virol. 2019, 93, doi:10.1128/JVI.01358-18.

11. Hartlage, A.S.; Cullen, J.M.; Kapoor, A. The Strange, Expanding World of Animal Hepaciviruses. Annu. Rev. Virol. 2016, 3, 53, doi:10.1146/ANNUREV-VIROLOGY-100114-055104.

12. Scheel, T.K.H.; Simmonds, P.; Kapoor, A. Surveying the Global Virome: Identification and Characterization of HCV-Related Animal Hepaciviruses. Antiviral Res. 2015, 115, 83–93, doi:10.1016/j.antiviral.2014.12.014.

13. Deinhardt, F.; Holmes, A.W.; Capps, R.B.; Popper, H. Studies on the Transmission of Human Viral Hepatitis to Marmoset Monkeys. I. Transmission of Disease, Serial Passages, and Description of Liver Lesions. J. Exp. Med. 1967, 125, 673–688, doi:10.1084/jem.125.4.673.

14. Stapleton, J.T.; Foung, S.; Muerhoff, A.S.; Bukh, J.; Simmonds, P. The GB Viruses: A Review and Proposed Classification of GBV-A, GBV-C (HGV), and GBV-D in Genus Pegivirus within the Family Flaviviridae. J. Gen. Virol. 2011, 92, 233–246, doi:10.1099/vir.0.027490-0.

15. Simons, J.N.; Pilot-Matias, T.J.; Leary, T.P.; Dawson, G.J.; Desai, S.M.; Schlauder, G.G.; Muerhoff, A.S.; Erker, J.C.; Buijk, S.L.; Chalmers, M.L.; et al. Identification of Two Flavivirus-like Genomes in the GB Hepatitis Agent (Hepatitis C Virus/Representational Difference Analysis); 1995; Vol. 92;.

16. Bukh, J.; Apgar, C.L.; Yanagi, M. Toward a Surrogate Model for Hepatitis C Virus: An Infectious Molecular Clone of the GB Virus-B Hepatitis Agent. Virology 1999, doi:10.1006/viro.1999.9941.

17. Dale, J.M.; Hood, S.P.; Bowen, O.; Bright, H.; Cutler, K.L.; Berry, N.; Almond, N.; Goldin, R.; Karayiannis, P.; Rose, N.J. Development of Hepatic Pathology in GBV-B-Infected Red-Bellied Tamarins (Saguinus Labiatus). J. Med. Virol. 2020, doi:10.1002/jmv.25769.

18. Li, T.; Zhu, S.; Shuai, L.; Xu, Y.; Yin, S.; Bian, Y.; Wang, Y.; Zuo, B.; Wang, W.; Zhao, S.; et al. Infection of Common Marmosets with Hepatitis C Virus/GB Virus-B Chimeras. Hepatology 2014, doi:10.1002/hep.26750.

19. Manickam, C.; Rajakumar, P.; Wachtman, L.; Kramer, J.A.; Martinot, A.J.; Varner, V.; Giavedoni, L.D.; Reeves, R.K. Acute Liver Damage Associated with Innate Immune Activation in a Small Nonhuman Primate Model of Hepacivirus Infection. J. Virol. 2016, 90, 9153–9162, doi:10.1128/JVI.01051-16.

20. Manickam, C.; Reeves, R.K. Modeling HCV Disease in Animals: Virology, Immunology and Pathogenesis of HCV and GBV-B Infections. Front. Microbiol. 2014, 5, 690, doi:10.3389/fmicb.2014.00690.

21. Marnata, C.; Saulnier, A.; Mompelat, D.; Krey, T.; Cohen, L.; Boukadida, C.; Warter, L.; Fresquet, J.; Vasiliauskaite, I.; Escriou, N.; et al. Determinants Involved in Hepatitis C Virus and GB Virus B Primate Host Restriction. J. Virol. 2015, 89, 12131–12144, doi:10.1128/JVI.01161-15.

22. Beames, B.; Chavez, D.; Guerra, B.; Notvall, L.; Brasky, K.M.; Lanford, R.E. Development of a Primary Tamarin Hepatocyte Culture System for GB Virus-B: A Surrogate Model for Hepatitis C Virus. J. Virol. 2000, 74, 11764–11772, doi:10.1128/JVI.74.24.11764-11772.2000.

23. Bukh, J.; Apgar, C.L.; Govindarajan, S.; Purcell, R.H. Host Range Studies of GB Virus-B Hepatitis Agent, the Closest Relative of Hepatitis C Virus, in New World Monkeys and Chimpanzees. J. Med. Virol. 2001, 65, 694–697, doi:10.1002/jmv.2092.

24. Lanford, R.E.; Chavez, D.; Notvall, L.; Brasky, K.M. Comparison of Tamarins and Marmosets as Hosts for GBV-B Infections and the Effect of Immunosuppression on Duration of Viremia. Virology 2003, 311, 72–80, doi:10.1016/S0042-6822(03)00193-4.

25. Schaluder, G.G.; Dawson, G.J.; Simons, J.N.; Pilot-Matias, T.J.; Gutierrez, R.A.; Heynen, C.A.; Knigge, M.F.; Kurpiewski, G.S.; Buijk, S.L.; Leary, T.P. Molecular and Serologic Analysis in the Transmission of the GB Hepatitis Agents. J. Med. Virol. 1995, 46, 81–90.

26. Sagan, S.M.; Sarnow, P.; Wilson, J.A. Modulation of GB Virus B RNA Abundance by MicroRNA-122: Dependence on and Escape from MicroRNA-122 Restriction. J. Virol. 2013, 87, 7338–7347, doi:10.1128/JVI.00378-13.

27. Lanford, R.E.; Chavez, D.; Guerra, B.; Lau, J.Y.N.; Hong, Z.; Brasky, K.M.; Beames, B. Ribavirin Induces Error-Prone Replication of GB Virus B in Primary Tamarin Hepatocytes. J. Virol. 2001, 75, 8074–8081, doi:10.1128/JVI.75.17.8074-8081.2001.

28. Bright, H.; Carroll, A.R.; Watts, P.A.; Fenton, R.J. Development of a GB Virus B Marmoset Model and Its Validation with a Novel Series of Hepatitis C Virus NS3 Protease Inhibitors. J. Virol. 2004, doi:10.1128/jvi.78.4.2062-2071.2004.

29. Pileri, P.; Uematsu, Y.; Campagnoli, S.; Galli, G.; Falugi, F.; Petracca, R.; Weiner, A.J.; Houghton, M.; Rosa, D.; Grandi, G.; et al. Binding of Hepatitis C Virus to CD81. Science (80-.). 1998, 282, 938–941, doi:10.1126/science.282.5390.938.

30. Scarselli, E.; Ansuini, H.; Cerino, R.; Roccasecca, R.M.; Acali, S.; Filocamo, G.; Traboni, C.; Nicosia, A.; Cortese, R.; Vitelli, A. The Human Scavenger Receptor Class B Type I Is a Novel Candidate Receptor for the Hepatitis C Virus. EMBO J. 2002, 21, 5017–5025, doi:10.1093/emboj/cdf529.

31. Evans, M.J.; von Hahn, T.; Tscherne, D.M.; Syder, A.J.; Panis, M.; Wölk, B.; Hatziioannou, T.; McKeating, J.A.; Bieniasz, P.D.; Rice, C.M. Claudin-1 Is a Hepatitis C Virus Co-Receptor Required for a Late Step in Entry. Nature 2007, 446, 801–805, doi:10.1038/nature05654.

32. Ploss, A.; Evans, M.J.; Gaysinskaya, V.A.; Panis, M.; You, H.; de Jong, Y.P.; Rice, C.M. Human Occludin Is a Hepatitis C Virus Entry Factor Required for Infection of Mouse Cells. Nature 2009, 457, 882–886, doi:10.1038/nature07684.

33. Baktash, Y.; Madhav, A.; Coller, K.E.; Randall, G. Single Particle Imaging of Polarized Hepatoma Organoids upon Hepatitis C Virus Infection Reveals an Ordered and Sequential Entry Process. Cell Host Microbe 2018, 23, 382-394.e5, doi:10.1016/j.chom.2018.02.005.

34. Owen, D.M.; Huang, H.; Ye, J.; Gale, M. Apolipoprotein E on Hepatitis C Virion Facilitates Infection through Interaction with Low Density Lipoprotein Receptor. Virology 2009, 394, 99, doi:10.1016/J.VIROL.2009.08.037.

35. Lupberger, J.; Zeisel, M.B.; Xiao, F.; Thumann, C.; Fofana, I.; Zona, L.; Davis, C.; Mee, C.J.; Turek, M.; Gorke, S.; et al. EGFR and EphA2 Are Host Factors for Hepatitis C Virus Entry and Possible Targets for Antiviral Therapy. Nat. Med. 2011, 17, 589, doi:10.1038/NM.2341.

36. Sainz, B.; Barretto, N.; Martin, D.N.; Hiraga, N.; Imamura, M.; Hussain, S.; Marsh, K.A.; Yu, X.; Chayama, K.; Alrefai, W.A.; et al. Identification of the Niemann-Pick C1-like 1 Cholesterol Absorption Receptor as a New Hepatitis C Virus Entry Factor. Nat. Med. 2012, 18, 281, doi:10.1038/NM.2581.

37. Wolfisberg, R.; Thorselius, C.E.; Salinas, E.; Elrod, E.; Trivedi, S.; Nielsen, L.; Fahnøe, U.; Kapoor, A.; Grakoui, A.; Rice, C.M.; et al. Neutralization and Receptor Use of Infectious Culture–Derived Rat Hepacivirus as a Model for HCV. Hepatology 2022, doi:10.1002/hep.32535.

38. Rey, F.A.; Lok, S.-M. Common Features of Enveloped Viruses and Implications for Immunogen Design for Next-Generation Vaccines. Cell 2018, 172, 1319–1334, doi:10.1016/j.cell.2018.02.054.

39. Hsu, M.; Zhang, J.; Flint, M.; Logvinoff, C.; Cheng-Mayer, C.; Rice, C.M.; McKeating, J.A. Hepatitis C Virus Glycoproteins Mediate PH-Dependent Cell Entry of Pseudotyped Retroviral Particles. Proc. Natl. Acad. Sci. U. S. A. 2003, 100, 7271–7276, doi:10.1073/pnas.0832180100.

40. Bartosch, B.; Dubuisson, J.; Cosset, F.-L. Infectious Hepatitis C Virus Pseudo-Particles Containing Functional E1–E2 Envelope Protein Complexes. J. Exp. Med. 2003, 197, 633–642, doi:10.1084/jem.20021756.

41. Drummer, H.E.; Maerz, A.; Poumbourios, P. Cell Surface Expression of Functional Hepatitis C Virus E1 and E2 Glycoproteins. FEBS Lett. 2003, doi:10.1016/S0014-5793(03)00635-5.

42. Tarr, A.W.; Urbanowicz, R.A.; Hamed, M.R.; Albecka, A.; McClure, C.P.; Brown, R.J.P.; Irving, W.L.; Dubuisson, J.; Ball, J.K. Hepatitis C Patient-Derived Glycoproteins Exhibit Marked Differences in Susceptibility to Serum Neutralizing Antibodies: Genetic Subtype Defines Antigenic but Not Neutralization Serotype. J. Virol. 2011, 85, 4246–4257, doi:10.1128/JVI.01332-10.

43. Logvinoff, C.; Major, M.E.; Oldach, D.; Heyward, S.; Talal, A.; Balfe, P.; Feinstone, S.M.; Alter, H.; Rice, C.M.; McKeating, J.A. Neutralizing Antibody Response during Acute and Chronic Hepatitis C Virus Infection. Proc. Natl. Acad. Sci. U. S. A. 2004, 101, 10149–10154, doi:10.1073/pnas.0403519101.

44. Bartosch, B.; Bukh, J.; Meunier, J.C.; Granier, C.; Engle, R.E.; Blackwelder, W.C.; Emerson, S.U.; Cosset, F.L.; Purcell, R.H. In Vitro Assay for Neutralizing Antibody to Hepatitis C Virus: Evidence for Broadly Conserved Neutralization Epitopes. Proc. Natl. Acad. Sci. U. S. A. 2003, 100, 14199–14204, doi:10.1073/pnas.2335981100.

45. Yamamoto, S.; Fukuhara, T.; Ono, C.; Uemura, K.; Kawachi, Y.; Shiokawa, M.; Mori, H.; Wada, M.; Shima, R.; Okamoto, T.; et al. Lipoprotein Receptors Redundantly Participate in Entry of Hepatitis C Virus. PLoS Pathog. 2016, 12, e1005610, doi:10.1371/journal.ppat.1005610.

46. Furuse, M.; Fujita, K.; Hiiragi, T.; Fujimoto, K.; Tsukita, S. Claudin-1 and -2: Novel Integral Membrane Proteins Localizing at Tight Junctions with No Sequence Similarity to Occludin. J. Cell Biol. 1998, 141, 1539–1550, doi:10.1083/jcb.141.7.1539.

47. Barton, E.S.; Forrest, J.C.; Connolly, J.L.; Chappell, J.D.; Liu, Y.; Schnell, F.J.; Nusrat, A.; Parkos, C.A.; Dermody, T.S. Junction Adhesion Molecule Is a Receptor for Reovirus. Cell 2001, 104, 441–451, doi:10.1016/S0092-8674(01)00231-8.

48. Bergelson, J.M.; Cunningham, J.A.; Droguett, G.; Kurt-Jones, E.A.; Krithivas, A.; Hong, J.S.; Horwitz, M.S.; Crowell, R.L.; Finberg, R.W. Isolation of a Common Receptor for Coxsackie B Viruses and Adenoviruses 2 and 5. Science (80-.). 1997, 275, 1320–1323, doi:10.1126/science.275.5304.1320.

49. Zeisel, M.B.; Dhawan, P.; Baumert, T.F. Tight Junction Proteins in Gastrointestinal and Liver Disease. Gut 2019, 68, 547–561, doi:10.1136/gutjnl-2018-316906.

50. Mee, C.J.; Harris, H.J.; Farquhar, M.J.; Wilson, G.; Reynolds, G.; Davis, C.; van IJzendoorn, S.C.D.; Balfe, P.; McKeating, J.A. Polarization Restricts Hepatitis C Virus Entry into HepG2 Hepatoma Cells. J. Virol. 2009, 83, 6211–6221, doi:10.1128/JVI.00246-09.

51. Mailly, L.; Xiao, F.; Lupberger, J.; Wilson, G.K.; Aubert, P.; Duong, F.H.T.; Calabrese, D.; Leboeuf, C.; Fofana, I.; Thumann, C.; et al. Clearance of Persistent Hepatitis C Virus Infection in Humanized Mice Using a Claudin-1-Targeting Monoclonal Antibody. Nat. Biotechnol. 2015, 33, 549–554, doi:10.1038/nbt.3179.

52. Harris, H.J.; Farquhar, M.J.; Mee, C.J.; Davis, C.; Reynolds, G.M.; Jennings, A.; Hu, K.; Yuan, F.; Deng, H.; Hubscher, S.G.; et al. CD81 and Claudin 1 Coreceptor Association: Role in Hepatitis C Virus Entry. J. Virol. 2008, 82, 5007–5020, doi:10.1128/jvi.02286-07.

53. Krieger, S.E.; Zeisel, M.B.; Davis, C.; Thumann, C.; Harris, H.J.; Schnober, E.K.; Mee, C.; Soulier, E.; Royer, C.; Lambotin, Melanie; et al. Inhibition of Hepatitis c Virus Infection by Anti-Claudin-1 Antibodies Is Mediated by Neutralization of E2-CD81-Claudin-1 Associations. Hepatology 2010, doi:10.1002/hep.23445.

54. Douam, F.; Dao Thi, V.L.; Maurin, G.; Fresquet, J.; Mompelat, D.; Zeisel, M.B.; Baumert, T.F.; Cosset, F.L.; Lavillette, D. Critical Interaction between E1 and E2 Glycoproteins Determines Binding and Fusion Properties of Hepatitis C Virus during Cell Entry. Hepatology 2014, 59, 776–788, doi:10.1002/hep.26733.

55. Hopcraft, S.E.; Evans, M.J. Selection of a Hepatitis C Virus with Altered Entry Factor Requirements Reveals a Genetic Interaction between the E1 Glycoprotein and Claudins. Hepatology 2015, 62, 1059–1069, doi:10.1002/hep.27815.

56. Haddad, J.G.; Rouillé, Y.; Hanoulle, X.; Descamps, V.; Hamze, M.; Dabboussi, F.; Baumert, T.F.; Duverlie, G.; Lavie, M.; Dubuisson, J. Identification of Novel Functions for Hepatitis C Virus Envelope Glycoprotein E1 in Virus Entry and Assembly. J. Virol. 2017, 91, doi:10.1128/jvi.00048-17.

57. Merz, A.; Long, G.; Hiet, M.-S.; Brügger, B.; Chlanda, P.; Andre, P.; Wieland, F.; Krijnse-Locker, J.; Bartenschlager, R. Biochemical and Morphological Properties of Hepatitis C Virus Particles and Determination of Their Lipidome. J. Biol. Chem. 2011, 286, 3018–3032, doi:10.1074/jbc.M110.175018.

58. Catanese, M.T.; Uryu, K.; Kopp, M.; Edwards, T.J.; Andrus, L.; Rice, W.J.; Silvestry, M.; Kuhn, R.J.; Rice, C.M. Ultrastructural Analysis of Hepatitis C Virus Particles. Proc. Natl. Acad. Sci. 2013, 110, 9505–9510, doi:10.1073/pnas.1307527110.

59. de la Peña, A.T.; Sliepen, K.; Eshun-Wilson, L.; Newby, M.; Allen, J.D.; Koekkoek, S.; Zon, I.; Chumbe, A.; Crispin, M.; Schinkel, J.; et al. Structure of the Hepatitis C Virus E1E2 Glycoprotein Complex. bioRxiv 2021, 2021.12.16.472992, doi:10.1101/2021.12.16.472992.

60. Kong, L.; Giang, E.; Nieusma, T.; Kadam, R.U.; Cogburn, K.E.; Hua, Y.; Dai, X.; Stanfield, R.L.; Burton, D.R.; Ward, A.B.; et al. Hepatitis C Virus E2 Envelope Glycoprotein Core Structure. Science 2013, 342, 1090–1094, doi:10.1126/science.1243876.

61. Khan, A.G.; Whidby, J.; Miller, M.T.; Scarborough, H.; Zatorski, A. V; Cygan, A.; Price, A.A.; Yost, S.A.; Bohannon, C.D.; Jacob, J.; et al. Structure of the Core Ectodomain of the Hepatitis C Virus Envelope Glycoprotein 2. Nature 2014, 509, 381–384, doi:10.1038/nature13117.

62. Flyak, A.I.; Ruiz, S.; Colbert, M.D.; Luong, T.; Crowe, J.E.; Bailey, J.R.; Bjorkman, P.J. HCV Broadly Neutralizing Antibodies Use a CDRH3 Disulfide Motif to Recognize an E2 Glycoprotein Site That Can Be Targeted for Vaccine Design. Cell Host Microbe 2018, 24, 703-716.e3, doi:10.1016/j.chom.2018.10.009.

63. Towers, G.; Bock, M.; Martin, S.; Takeuchi, Y.; Stoye, J.P.; Danos, O. A Conserved Mechanism of Retrovirus Restriction in Mammals. Proc. Natl. Acad. Sci. U. S. A. 2000, 97, 12295–12299, doi:10.1073/pnas.200286297.

64. Zufferey, R.; Nagy, D.; Mandel, R.J.; Naldini, L.; Trono, D. Multiply Attenuated Lentiviral Vector Achieves Efficient Gene Delivery in Vivo. Nat. Biotechnol. 1997, 15, 871–875, doi:10.1038/nbt0997-871.

65. Hitoshi, N.; Ken-ichi, Y.; Jun-ichi, M. Efficient Selection for High-Expression Transfectants with a Novel Eukaryotic Vector. Gene 1991, 108, 193–199, doi:10.1016/0378-1119(91)90434-D.

66. Kalemera, M.D.; Capella-Pujol, J.; Chumbe, A.; Underwood, A.; Bull, R.A.; Schinkel, J.; Sliepen, K.; Grove, J. Optimized Cell Systems for the Investigation of Hepatitis C Virus E1E2 Glycoproteins. J. Gen. Virol. 2021, 102, doi:10.1099/jgv.0.001512.

67. Heckman, K.L.; Pease, L.R. Gene Splicing and Mutagenesis by PCR-Driven Overlap Extension. Nat. Protoc. 2007, 2, 924–932, doi:10.1038/nprot.2007.132.

68. Fukuhara, T.; Kambara, H.; Shiokawa, M.; Ono, C.; Katoh, H.; Morita, E.; Okuzaki, D.; Maehara, Y.; Koike, K.; Matsuura, Y. Expression of MicroRNA MiR-122 Facilitates an Efficient Replication in Nonhepatic Cells upon Infection with Hepatitis C Virus. J. Virol. 2012, 86, 7918–7933, doi:10.1128/jvi.00567-12.

